# Hypergravity affects Cell Traction Forces of Fibroblasts

**DOI:** 10.1101/2020.03.30.015958

**Authors:** J. Eckert, J. J.W.A. van Loon, L. M. Eng, T. Schmidt

**Affiliations:** Leiden University; Amsterdam University; Dresden University of Technology

## Abstract

Cells sense and react on changes of the mechanical properties of their environment, and likewise respond to external mechanical stress applied to them. Whether the gravitational field, as overall body force, modulates cellular behavior is however unclear. Different studies demonstrated that micro- and hypergravity influences the shape and elasticity of cells, initiate cytoskeleton reorganization, and influence cell motility. All these cellular properties are interconnected, and contribute to forces that cells apply on their surrounding microenvironment. Yet, studies that investigated changes of cell traction forces under hypergravity conditions are scarce. Here we performed hypergravity experiments on 3T3 fibroblast cells using the Large Diameter Centrifuge at the European Space and Technology Centre (ESA-ESTEC). cells were exposed to hypergravity of up to 19.5*g* for 16 h in both the upright and the inverted orientation with respect to the g-force vector. We observed a decrease in cellular traction forces when the gravitational field was increased up to 5.4*g*, followed by an increase of traction forces for higher gravity fields up to 19.5*g* independent of the orientation of the gravity vector. We attribute the switch in cellular response to shear-thinning at low g-forces, followed by significant rearrangement and enforcement of the cytoskeleton at high g-forces.

**SIGNIFICANCE:** The behavior of cells critically depend on the mechanical properties of their environment. For example external stresses and strains lead to decisions in cell differentiation as well as to collective-migration in metastasis. Gravity, as a permanently acting body force, is one of those externs stresses. We demonstrate the impact of gravitational challenges on forces that cells apply to their environment. We observed a switch in cellular response with a decrease in cell traction forces for low bypgrarayiv. conditions, followed by a significant increase in cell traction forces at higher g-level. This particular cellular response reflects a switch in croskeletal organization, similar to that observed for cells in fluids where shear forces act.

## INTRODUCTION

In recent years it became accepted that cellular function is, in part, controlled by external mechanical cues. Mechanical cues were shown to be sufficient to differentiate mesenchymal stem cells (1), initiate transcriptional programs (2), drive morphogenesis (3), direct cell migration (4), and control malignancy in tumors (5). The force and stiffness-mediated responses of cells on the mechanical properties of the extracellular matrix (ECM) are attributed to, yet to be identified, mechano-chemical sensor platforms that transform external mechanical cues into intracellular biochemical signals, ultimately leading to e.g. altered gene expression (6). The multi-protein sensory units responsible for mechano-sensation are summarized as cell-matrix adhesions, focal adhesions, and cell-cell adhesions (7). In focal adhesions, transmembrane receptor proteins such as integrins (8), bind to specific proteins of the ECM. On the cytosolic side those proteins link, through a extended protein-cascade, to the actin cytoskeleton, the cellular machinery that can apply forces through the contraction of actin fibers through the linking myosin motor-activity (9–11).

So far, studies on the mechano-chemical coupling have focused on cellular responses related to static extracellular stiffness (12) and topography (13), as well as on the direct mechanical stimulation of cells by fluid flow (14), micropipette aspiration (15) (16), optical tweezers (17), optical stretchers (18), atomic-force microscopes (19), and magnetically actuated particles (20). In most of those experiments local stress was applied to the cells that resulted in a highly sensitive cellular response. Cells were shown to adapt to their local mechanical environment by enforcing their cytoskeleton. Such restructuring of the cytoskeleton is paralleled by an increased force application of cells when placed into stiffer environments or when challenged by higher external tensions.

Surprisingly, the robust cellular response on localized mechanical cues appears more subtle for homogeneous mechanical cues such as that given by gravity. Experiments showed that hypergravity in the range of 2-20*g* does influence cellular morphology and elasticity (21) (22), the cytoskeletal organization (23) and the motility (24). The strength and impact, however, differ largely, as was reviewed for endothelial cells by Maier *et al.* (25). The effects observed depend on many factors, which appeared difficult to disentangle. Next to different cell types, different g-levels applied, and different exposure times of challenge, in particular, the use of small laboratory centrifuges by which hypergravity was produced are known to generate additional mechanical cues on cells (in particular internal shear-forces) that possibly overshadow the effects of gravitational cues (26).

Monici *et al.* exposed microvascular endothelial cells for a period of 5 × 10 min to 10g. Cells were centrifuged under closed conditions in a thermostated laboratory centrifuge. In the experiment, the authors showed changes in the cytoskeleton organization (23). The opposite was reported by Costa-Almeida *et al* (27). In the latter experiment endothelial cells (HUVEC) were exposed exposed to 3 and 10g for 4 and 16 h using the large-diameter centrifuge (LDC) at the ESA/ESTEC center in Noordwijk. No changes in cytoskeleton organization was observed.

Here, we focus on investigating the impact of hypergravity as a body force on the active cellular mechano-response. Given that hypergravity has an influence on the cell morphology (21) (22), cytoskeleton organization (23), membrane viscosity (28), and motility (24), we probed whether or not cells react on hypergravity by a modulation of their traction forces towards the ECM.

## MATERIALS AND METHODS

### Cell Culture

3T3 fibroblasts were cultured in high-glucose Dulbecco’s Modified Eagle’s Medium (DMEM, Sigma Aldrich) supplemented with 10% fetal calf serum (FCS, Thermo Fisher Scientific), 2mM glutamine and 100 μg/ml penicillin/streptomycin, 37°C, 5% CO_2_.

### Immunostaining

5 min after the hypergravity exposure, cells were fixed for 15 min in 4% paraformaldehyde (Alfa Aesar, 43368) in phosphate-buffered saline (PBS). After fixation, cells were permeabilized for 10min with 0.1% Triton-X and blocked for 60min with 1% BSA in PBS. F-actin was stained with Alexa Fluor 532-labelled phalloidin (Invitrogen, A22282) and the DNA with DAPI (Sigma).

### Hypergravity Exposure

In order to avoid shear forces (26), hypergravity experiments were performed using the Large Diameter Centrifuge (LDC, Fig. 1A) at the European Space Research and Technology Center in Noordwijk, The Netherlands. Cells were seeded on two sets of elastic micropillar arrays, located in 12-well-plates with one array per well, and incubated for 5.5 h, 37°C, 5% CO_2_. After cell spreading on top of the functionalized micropillars, arrays of one set were flipped in the up-side-down orientation. Both sets were placed in closed metal boxes flushed with 5% CO_2_ at 100% humidity and stored inside the incubators of the LDC-gondolas held at 35 - 37°C (Fig. 1B). The centrifuge gondolas were placed at a distance of 2 and 4 m to the centrifuge axis, which allowed us to address two g-levels, simultaneously (29). 6.5 h after cell seeding, they were exposed for 16 h to hypergravity of 5.4*g*, 10*g* and 19.5*g*, respectively, as controlled by the distance of the gondolas from the axis, and the speed at which the centrifuge turned. As of the large-diameter of the centrifuge, shear-forces were negligible in our experiments (26). The g-force acted perpendicular to the sample surface (Fig. 1C). 1*g* control experiments were prepared and conducted simultaneously under identical conditions outside the centrifuge.

**Figure 1:**
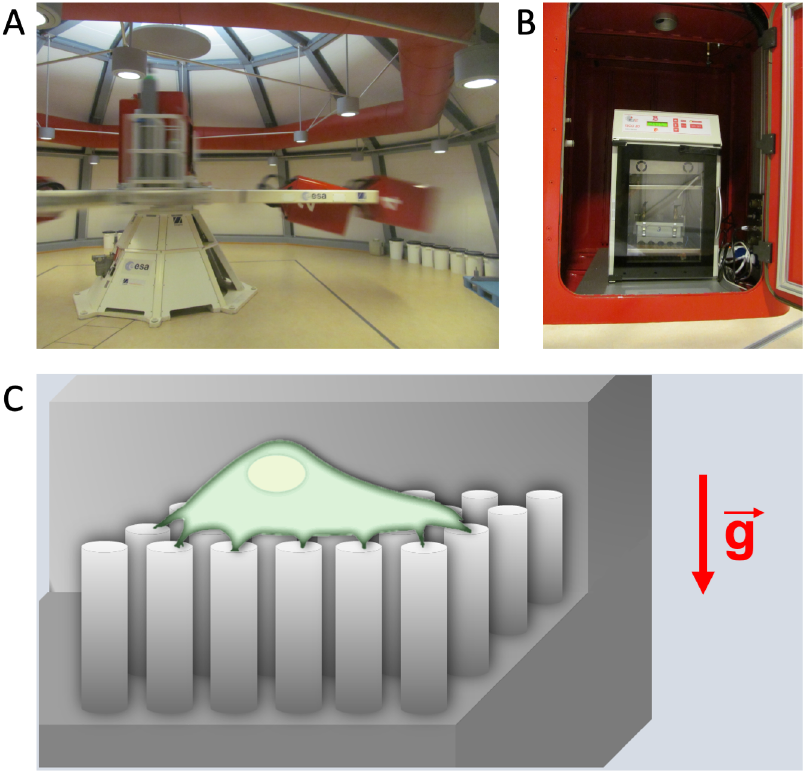
A: Large Diameter Centrifuge (LDC) at ESA/ESTEC provided hypergravity conditions. B: Samples were located in a metal box flushed with 5% CO_2_ inside the incubator of on one of the LDC-gondolas. The temperatur was set to 37°C. C: Schematic illustration of a cell assembled on top of a micropillar array in upright sample orientation. The g-force was exerted perpendicular to the cell spreading area.

### Elastic Micropillar Arrays

Polydimethylsiloxane (PDMS, Sylgard 184) micropillar arrays of 2 μm diameter, 6.9 μm height and 4 μm spacing in a hexagonal geometry were used for cell traction force experiments. The pillar arrays were flanked by 50 μm spacers on two sides of the array. Details of this arrangement and the experimental procedures was described earlier in detail (30). In brief, pillar arrays were produced on a negative silicon-wafer master made by a two-step deep reactive-ion etching process. Wafers were passivated in trichloro-silane (Sigma Aldrich, 448931). A mixture of 1:10 PDMS (crosslinker:base ratio) was poured onto the Si-master and cured for 20 h at 110°C. After peel-off, the tops of the pillars were coated by μ-contact printing. For that, flat 1:30 PDMS stamps were incubated for 1 h with 40 μl of 50 μg/ml Alexa Fluor 647 labelled and 50 μg/ml unlabelled fibronectin (Sigma Aldrich, F1141), then washed and dried. Subsequently, the stamps were gently loaded onto the UV-ozone-activated micropillar arrays for 10 min. After stamping, the arrays were passivated with 0.2% Pluronic (F-127, Sigma Aldrich, P2443) for 1 h, and washed in phosphate-buffered saline (PBS).

### Microscopy

Samples were imaged at high resolution on a home-build optical microscope setup based on an inverted Axiovert200 microscope body (Zeiss), a spinning disk unit (CSU-X1, Yokogawa), and an emCCD camera (iXon 897, Andor). IQ-software (Andor) was used for setup-control and data acquisition. Illumination was performed using fiber-coupling of different lasers (405 nm (CrystalLaser), 514nm (Cobold) and 642nm (Spectra-Physics Excelsior)). Pillar arrays were placed up-side-down onto 25 mm cover glasses and inspected with an EC Plan-NEOFLUAR 40×1.3 Oil immersion objective (Zeiss).

### Image Analysis

Images were analysed using Matlab scripts (Mathworks, Matlab R2017a). Pillar deflections were quantified as previously described in detail (30). The accuracy of the analysis was determined from an undeflected area of the pillar array by selecting a pillar region outside the cell area. Pillar deflections underneath the cell within the background range were discarded. The traction force per pillar was calculated by dividing the total absolute force per cell by the number of deflected pillars per cell. The cell spreading area was calculated as the number of deflected pillars per cell multiplied by the unit-cell area of the hexagonal pillar array geometry. The unit-cell area measured 13.84 μm^2^.

Additionally, we calculated the bending modulus of pillars caused by the increase in the weight of the cell and the pillar itself at higher g-level (see supplemental material). Based on the study by Grover *et al.*, we assumed a cell density of 1.08 g ml^-1^ (31). The averaged diameter of 3T3 fibroblasts is 50 μm with a height of 15 μm measured via z-stack images. Assuming a deflection of 400 nm, the differences of the pillar deflection at 1g to that at 20*g* is 4.7 pN (S5). Hence, the additional pillar deflection is 3 orders of magnetude lower than that resulting from cellular traction force, and can thus be neglected.

### Statistics

The following samples with cells were analyzed in the upright orientation: Two control arrays with 101 cells at 1*g*, one array with 72 cells at 5.4*g*, two arrays with 112 cells including one repeat at 10*g*, and one array with 66 cells at 19.5*g*.

In the up-side-down orientation, we analyzed: Two control arrays with 100 cells at 1*g*, one array with 65 cells at 5.4*g*, one array with 54 cells at 10*g*, and one array with 47 cells at 19.5*g*. Hence, in total 617 cells and 20343 deflections were analyzed.

P-values were calculated using the two-sided Wilcoxon rank sum test in Matlab. Data sets were significantly different with probabilities of p≤0.05 (*); p<0.01 (**); p<0.001 (***).

## RESULTS

Given the results of prior studies on the effect of hypergravity on cellular behavior (21) (23) (24), we anticipated that the effect of hypergravity on cellular force application would be small. Hence, we initially performed an extensive analysis of cell traction forces using the micropillar technique to extract a solid, experimentally confirmed base-line value for all parameters investigated.

Cells were seeded on six independently produced arrays and left for 6 h. After fixation and staining for actin and DNA, 323 cells leading to 9545 pillar deflections were analyzed. In a first step, the total absolute force that a cell produces was compared to the number of deflected pillars for that particular cell. The data are shown in Fig. 2A. The total force per cell appeared highly correlated to the number of deflected pillars (|*r*| > 0.8). From this correlation, we concluded that the mean force per pillar is a robust descriptor for cellular force application. This conclusion corroborates earlier experiments in which it was shown that the force per pillar is a cellular property which depends on external chemical or mechanical cues (32). Hence, in what follows, we calculated the mean traction forces per deflected pillar.

**Figure 2:**
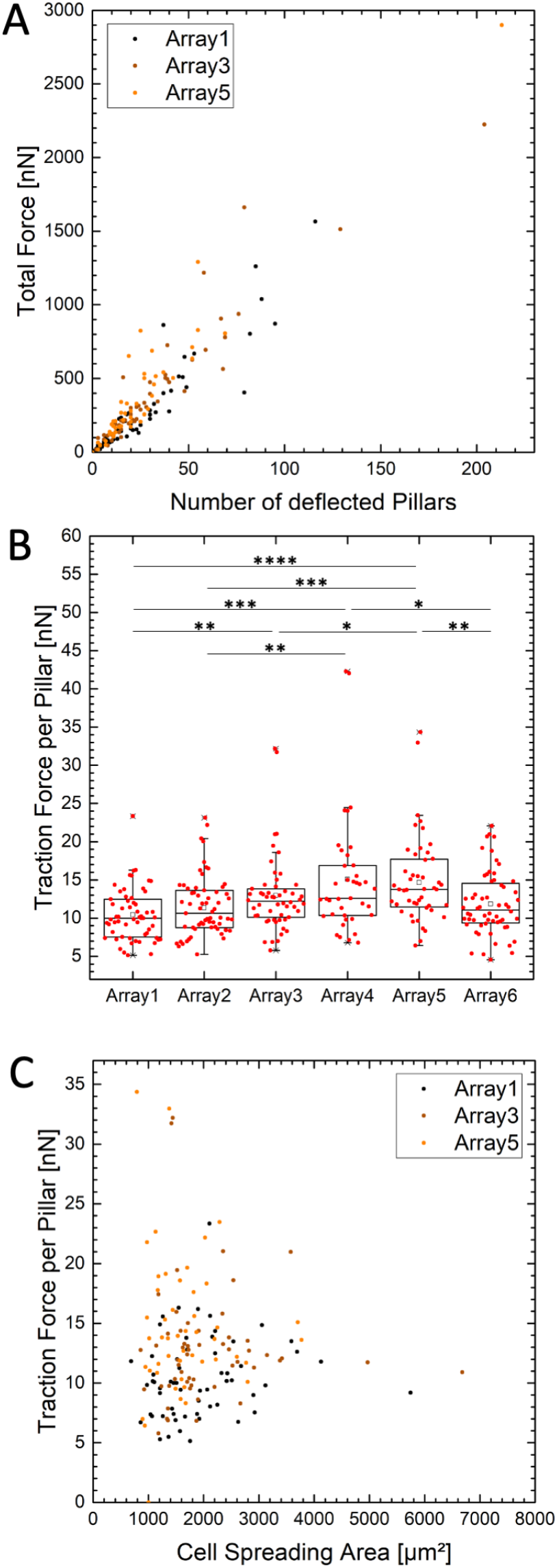
3T3 fibroblasts on six independently produced array under standard gravity conditions were analysed. The total force per cell increases linearly with the number of deflected pillars (A). The ratio results in the traction force per deflected pillar (B). This value differes significantly for cells on equally prepared arrays. The traction force is independent from the cell spreading area (C). Three of six datasets randomly chosen are shown in A and C. Each datapoint represent one analysed cell.

In how far the mean traction force per pillar is a robust quantity with respect to the production of micropillar arrays, we compared data from six independently produced arrays (Fig. 2B). Mean values for the force per pillar varied for each array between (10.4 ± 3.4) nN (array 1, 59 cells) and (15.1 ± 7.8) nN (array 4, 39 cells). A two-sided Wilcoxon rank sum test confirmed that the data set of each of the array was different. Hence, this result suggests that biological and technological (e.g. pillar geometry) variability in our data could easily be underestimated. In what follows, we thus defined results of cellular forces that fall into the range 10.4 −15.1 nN as indistinguishable from the control experiment at 1g-condition.

Further, we compared the cell size to the cellular force per pillar (Fig. 2C). The cell size was calculated from the cell spreading area as seen in the actin-channel of the images taken. In contrast to the strong correlation between total cell force and number of deflected pillars in Fig. 2A, the correlation coefficient between force per pillar and cell size was small, 0.05 < |*r*| ≤ 0.27. Hence, taken both results into account we concluded, that the mean force per deflected pillar is a robust measure of cellular forces when technological variability is properly considered.

Subsequently, we compared the 1*g* results to those obtained for hypergravity conditions. Cell-loaded micropillar arrays immersed in 12-well-plates were placed into the incubators of two gondolas of the large diameter centrifuge (LDC) at ESA/ESTEC. Together with the control at standard 1*g* gravity condition, we exposed cells to three different g-levels. For statistical reasons, we performed two independent experiments for 16h each at 1*g*, 5.4*g* and 10*g* and one experiment at 1*g*, 10*g* and 19.5*g*, respectively. The g-vector acted perpendicular to the cell spreading area. Arrays were located in both the upright (positive g-vector) and the up-side-down (negative g-vector) orientation. For analysis, cells were fixed, stained and imaged remotely.

As predicted from our 1*g* experiments, the total force per cell highly correlated with the number of deflected pillars also for hypergravity conditions for both the upright and the upside-down orientation. Data for the upright orientation (positive g-vector) are shown in Fig. 3A. The correlation coefficient of 0.8 ≤ |*r*| ≤ 1 was equivalent to that we found for 1*g*. In addition, we verified that the force per pillar was independent on the cell spreading area. The correlation is shown in Fig. 3D for two sets at two different g-levels. The data were uncorrelated as inferred from the low value of the correlation coefficient, |*r*| < 0.11. Hence, as for the 1*g* condition, the mean force per deflected pillar is a robust measure to characterize cellular forces also for hypergravity conditions.

**Figure 3:**
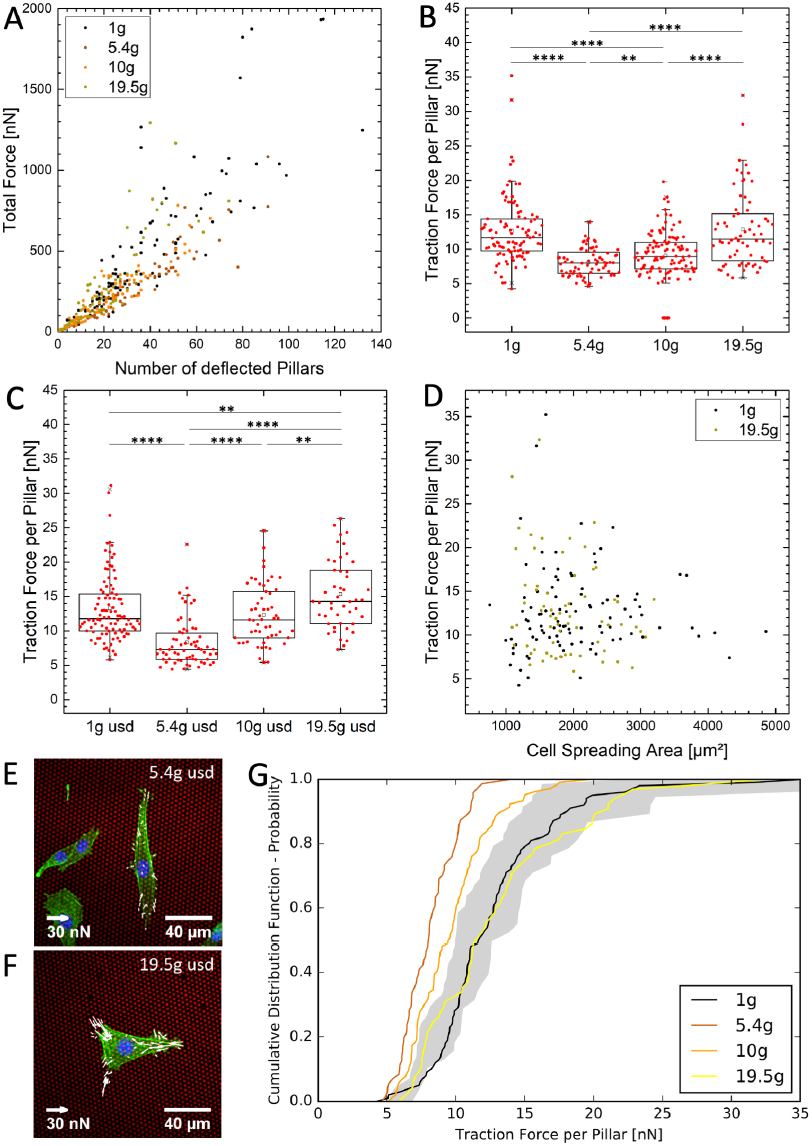
3T3 firoblasts were exposed to hypergravity in upright and up-side-down orientation. The total force per cell increases linearly with the number of deflected pillars of cells exposed to 1*g*, 5.4*g*, 10*g* and 19.5*g* in upright orientation (A). The ratio results in the traction force per deflected pillar for cells in upright (B) and up-side-down (usd) orientation (C). The traction force decreases from 1*g* to 5.4*g* and increases from 5.4*g* to 19.5*g*. This effect is independent from the cell spreading area (D). Two of four randomly chosen datasets are shown. Each data point represent one analysed cell. In the usd orientation, the biggest difference is between 5.4*g* (E) and 19.5*g* (F). Red: pillar array, green: actin filament, blue: nucleus. Compared to experiments under 1*g* gravity conditions, cells in upright orientation exert less force on pillars at 5.4*g* and 10*g* (G). The gray range is the defined force limit for cell traction forces under 1*g* conditions. This limit is set by the minimal and maximal averaged cell force of the six micropillars at 1*g* in Fig. 2B.

In turn, we analyzed the averaged force per deflected pillar for the various hypergravity levels (Fig. 3B-C). In both orientations, the force per pillar decreased significantly when hypergravity was changed from 1*g* to 5.4*g*. When hypergravity was increased further to 10*g* and 19.5*g*, respectively, the force per pillar increased again, finally exceeding the value at 1*g*. In the upright orientation (positive g-vector, Fig. 3B), cells applied significantly less force on pillars at 5.4*g* with an average of (8.1 ± 2.0) nN and (9.1 ± 3.7) nN at 10*g*, respectively. At 19.5*g*, the traction force per deflected pillar with an average of (12.9 ± 5.4) nN was not significantly different from 1*g* with (12.7 ± 4.8) nN. The strongest effect and biggest difference was measured for cells in up-side-down orientation (negative g-vector, Fig. 3C). In this orientation, cells applied the lowest force per pillar at 5.4*g* with an average of (8.3 ± 3.5) nN and the highest force with (15.3 ± 5.0) nN at 19.5*g*, which is visualized in Fig. 3E-F. The traction force of the 1*g* control with (13.3 ± 4.9) nN was significantly different to both results at 5.4*g* and 19.5*g*, while it was equivalent to the averaged force of (12.3 ± 4.4) nN at 10*g*. A two-sided Wilcoxon rank sum test was used to compare the two orientations shown in Fig. 3B-C. The force per pillar did not differ significantly for the 1*g* and the 5.4*g* situations. Yet, at higher g-levels the two configurations were significantly different, with *p* < 0.0001 at 10*g*, and *p* = 0.0044 for 19.5*g*.

A clearer picture of the significance of the different results was obtained from the analysis of the cumulative distribution function of the mean force per pillar values. From the six 1 *g*-control measurements, we constructed a significance-band for the cumulative distribution (gray area in Fig. 3G). The band was constructed such that all six 1g-control measurements fall into the given range. The forces of cells in the upright orientation exposed to 5.4*g* and 10*g* are fully outside of the gray range, while 1*g* and 19.5*g* samples are equal to the standard gravity distributions. This confirms our results of the change of cell traction forces under hypergravity conditions when analyzed by a Wilcoxon test (Fig. 3B).

## DISCUSSION

It has been shown that hypergravity influences cells morphology and behavior. Although the outcome may highly vary (25) it was well documented that hypergravity leads to a rearrangement of the cytoskeleton (23)(27). Here, we investigated in how far hypergravity modulates the contractile behavior of cells. Given that the cytoskeleton represents the main contractile machinery within cells, we anticipated that forces exerted by cells onto their environment, would likewise change.

We cultured 3T3 fibroblasts on elastic micropillar arrays and exposed them for 16h to hypergravity at a range of g-levels. As we predicted, cellular forces changed for hypergravity conditions: we found an initial decrease of forces from 1*g* to 5.4*g*, followed by a subsequent increase in forces at least up to 19.5*g*.

Our results corroborate earlier findings elucidating the reorganization of the actin network. Versari *et al.* cultured epithelial cells (HUVEC) in a medium-sized centrifuge for acceleration research (MidiCAR) and found less dense actin fibers after 96 h at 3.5*g* (33). The decrease in actin stress fibers at low hypergravity levels confers with our results on a decrease in traction forces at 5.4*g*. Also the more recent results by Costa-Almeida *et al*, who exposed human tendon-derived cells (hTDCs) to 5*g*, 10*g*, 15*g* and 20*g* for 4h and 16h (34), align well with our interpretation. After 4h, the anisotropy of actin fibers in hTDCs was significantly lower at 5*g* as compared to 1*g* and increased towards higher g-levels.

Other external stress measurements showed a similar effect on the actin filament formation. Kataoka *et al.* performed flow-imposed experiments. Using a parallel flow plate chamber, endothelial cells were exposed to different flow directions for 24 h. Under fluid shear stress as low as 2 Pa, cells perfectly aligned with the flow direction and formed thick stress fibers (35). Kuo *et al.* applied an oscillatory shear stress with a frequency of 1 Hz on cells. Under lower shear stress, the phalloidin-labelled F-actin signal decreases at 0.05 Pa (36) after 0.5 h, which indicates a reorganization of actin filaments. In terms of the cell volume (density: 1.08 g ml^-1^ (31), height: 15 μm), 1*g* can be assigned to a shear stress of 0.16 Pa, 10*g* to 1.6 Pa and 20*g* to 3.2 Pa. Hence, fluid-shear stress and hypergravity seem to have a similar impact on the actin reorganization of cells and possibly on cell traction forces. On the other hand, Perrault *et al.* measured an increase in strain energy independent from the flow rate (0.014 – 0.133 Pa) (37). Care must be taken that fluid-shear stress as a surface force acts parallel to the spread cell and only on the affected surface area. In contrast, hypergravity as a body force acts perpendicular to it and on all parts of the cell volume.

Further it was found that the internal organization of the cytoskeleton alters with mechanical challenge. Norstrom *et al.* studied shear thickening of cross-linked F-actin networks. Performing rheological experiments, the authors observed viscous deformation of stresses from 0.001 to 10 Pa, caused by stress stiffening and shear thickening. To a surprise, and in contrast to earlier findings in which stress weakening of sparsely cross-linked actin network was measured, Norstrom *et al.* observed a stress stiffening behaviour of a densely cross-linked network (38). This finding is consistent with the work of Gardel *et al.* (39) who highlighted the connection between the elasticity of the actin network, the density of cross-linkers and the actin concentration.

Combining the data of Norstrom *et al.* and Gardel *et al.* with our findings, an interaction between the external g-level acting on the cell and the elasticity of it’s actin network appears apparent. At low hypergravity levels, a less dense cross-linked actin network with less oriented stress fibers might be formed. This would result in a decrease in cell traction forces. At larger g-levels, the actin network would then form oriented densely-packed stress fibers, which are highly cross-linked. Those densely-packed, highly cross-linked stress fibers will result in higher traction forces due to stress stiffening.

Hence, based on our results, we propose that hypergravity causes a re-organization of the actin network dependent on the g-level, where gravitation acts similar to that reported for fluid-shear stress. Reorganization of the cytoskeleton subsequently causes a change in traction force that was observed in our experiments.

## CONCLUSION

In conclusion we demonstrated that hypergravity modulates the traction force of 3T3 fibroblasts. Dependent on the g-force level, the cell traction force first decreases for low hypergravity conditions, yet increases for higher g-levels. We found that cells in up-side-down orientation were more affected when compared to cells in upright orientation. We propose that the change in cellular force-response reflects the reorganization of the cytoskeleton as triggered by a gravitational cue, very similar to cellular responses to fluid shear-flow. Further studies should be employed to investigate the involvement of e.g. the myosin activity, and the actin stress fiber formation on the force transduction at altered gravity conditions. Our data should be considered in order to estimate potential health-riscs in planned long-haul space flights.

## AUTHOR CONTRIBUTIONS

sample preparation: JE

conducted hypergravity experiments: JE, JJWAL

data analysis: JE, TS

writing manuscript: JE, JJWAL, LME, TS

initiated the study: JJWAL

All authors gave final approval for publication.

## ACKNOWLEDGMENTS

We acknowledge Robert Lindner and Alan Dowson (TEC-MMG, ESA/ESTEC) for the support and access to the LIS-Lab and the LDC. J.E. thanks for the support of an Erasmus+ fellowship.

## SUPPLEMENTARY MATERIAL

### Beam Theory under Hypergravity Conditions

The deflection of the micropillar as an elastostatic process is described by the Bernoulli beam theory. Perpendicular acting forces to the beam axis with small angular changes are described by the Euler-Bernoulli assumption for narrow beams. The deflection of the beam *δ*:= *ω(x)* is calculated from the curvature of the bending line as a differential equation:

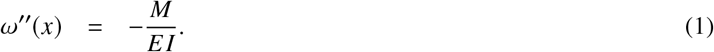

*M* is the bending moment, *E* the Young’s modulus and *I* the second momentum of area:

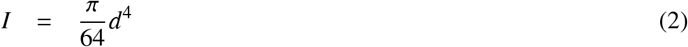

including the beam diameter, d. Taking the boundary conditions for a beam with clamping ω(0) = 0 and ω’(0) = 0, the lateral force acting on the pillar is described by

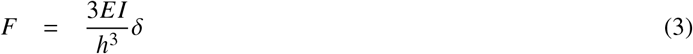

with the pillar height.

Gravity causes additional forces acting on the beam (Fig. 4A): the gravitational force of the pillar, *F*g*,_Pillar_*, and of the cell, *F*g*,_cell_*, both are calculated by the differential Eq. (1) with the moment of *M* = *F*g*z*. As a superposition, for the bending of the pillar it results:

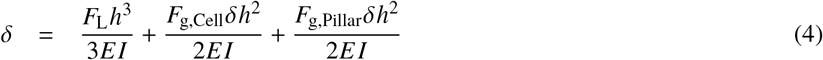

and the total acting force

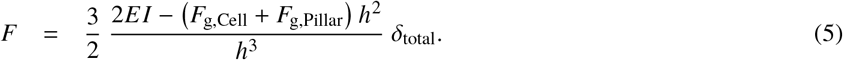

**Figure 4:**
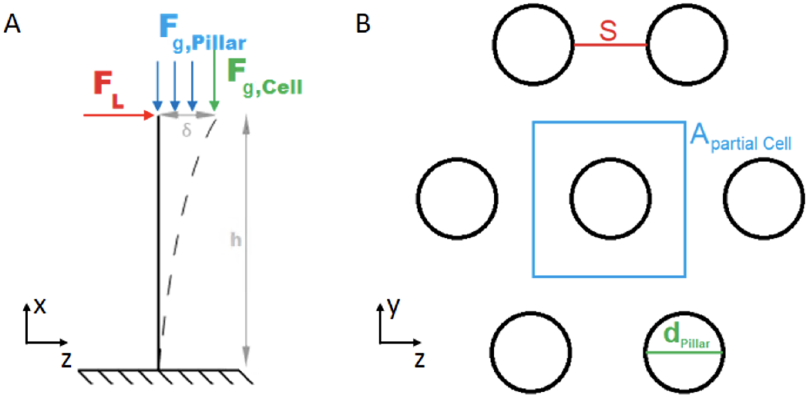
A: Micropillar is described by a cantilevered beam with the height, *h*. A cell sitting on the tip produces a lateral force, *F*, and causes a deflection, δ, of the pillar. In addition, gravitational forces increase the weight of the cell, *F_g,cell_*, and the pillar, *F_g,Pillar_*, causing a higher deflection. B: Schematic top view of the hexagonal micropillar array. *S* is the spacing, *d_Pillar_* the diameter of the pillar and *A_partial-Cell_* the partial area of the cell that acts on one single pillar

The gravitational forces are calculated by

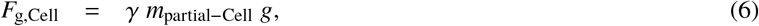

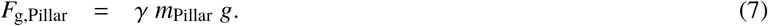

Here, *γ* is the gravitational level, g the gravitational acceleration, *m_Pillar_* the mass of one pillar and m_partial-Cell_ the partial mass of the cell that acts on one pillar. This partial mass can be calculated by using the ratio of the volumes:

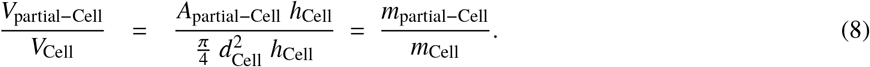

The height of the cell, *h_Cell_*, with a cylindrical volume cancels out. Thus, the partial mass of the cell only depends on three cell values: the mass, *m_Cell_*, the diameter, *d_Cell_*, and the partial area, *A_partial-Cell_*. The partial area is the part of the cell that acts on one pillar and can be calculated with the geometry of a hexagonal structure (Fig. 4B):

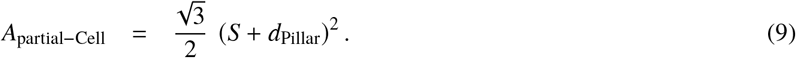

Here, *S* is the spacing between two pillars and d_Pillar_ the diameter of the pillar. The substitution of (9) into (8) yields the mass of the partial cell in (6):

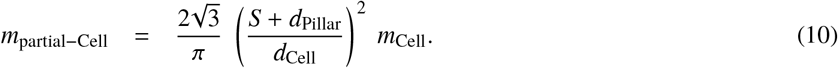

The mass of the pillar in (7) is given by

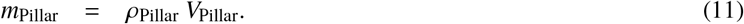

Here, *ρ*_Pillar_ is the density of the pillar material and *V_Pillar_* its cylindrical volume.

